# Developing tree improvement strategies for challenging environmental stresses under global climate change: a review from traditional tree breeding to genomics of adaptive traits for the quaking aspen

**DOI:** 10.1101/2023.08.25.554698

**Authors:** Deyu Mu, Chen Ding, Hao Chen, Yang Li, Earl M. (Fred) Raley

**Author notes:** Correspondence: CD. CD, DM, and HC contributed equally to this paper.

## Abstract

Quaking or trembling aspen in North America and Euro-Asia *(Populus tremuloides* and *P. tremula,* respectively) are both widely distributed species with a long history of scientific research and tree improvement work in areas such as carbon sequestration, biomass, bioenergy, wood, and fiber, as well as studies evaluating the social, economic, and ecological benefits of the species. This chapter reviews the ecological genetics and genomics of quaking aspen’s adaptive traits with a broad perspective of the relationship between phenotypic variation and genetic (G) and environmental (E) effects as well as their interactions (GxE). Based on recent studies, several adaptive traits are discussed, including spring and fall phenology and stress tolerance to environmental factors such as frost, salinity, drought, heat, UV radiation, etc. We also conducted a meta-analysis of empirical studies on adaptive traits of *P. tremuloides* and its sister species, as research using *P. tremuloides* as a true “model species” is currently limited. However, molecular tools and experimental designs in the form of different common gardens constitute an integrated pathway for the development of traits and varieties/populations to promote reforestation under changing climatic conditions.

## 1. Introduction

*Populus* trees are important broadleaf species for global forest ecosystems and for humans because they serve as model species for forest genetics, genomics studies, as well as tree improvement (Stanton et al., 2010). Quaking aspen (*Populus tremuloides* Michx.) and common or trembling aspen (*Populus tremula* L.) from the section *Populus* are widespread in boreal forests of North America and Eurasia and are of great economic and ecological importance (Barnes, 1975; Stettler et al., 1996), spanning a range of 110° ∼ 155° longitude and 50° ∼ 55° latitude on the two continents (Perala, 1990; Stanton et al., 2010). *P. tremuloides* has the largest natural range among native tree species in North America (Dickmann, 2002) and grows in various types of habitats that provide important resources for wildlife and plants (Canadian Forest Service, 2012; Stelfox, 1995). Due to the sexual and asexual reproductive modes in natural stands, *P. tremuloides* can occur and persist over long periods of time, not to mention sites where aspen may occur that are prone to abiotic stresses such as drought (Blonder et al., 2019; Mock et al., 2012). Severe drought and disturbances are causing increasing maladaptation and mortality of natural aspen stands in both the US and Canada (Ding et al., 2017; Rogers et al., 2020; Worrall et al., 2013). Globally, quaking aspen forests are projected to decline across landscapes due to increasing climate-related factors (i.e., heat and drought), as well as disturbance from pests, pathogens, and severe wildfires (Anderegg et al., 2013; Marchetti et al., 2011; Worrall et al., 2013). Rehfeldt et al. (2001) suggested that an evaluated redistribution of trees and populations across the species’ range could potentially improve adaptation to future climate stresses. Plausible prescriptions that include assisted migration will maintain and advance the productivity and health of natural and plantation forests. *P. tremuloides* will benefit from assisted migration strategies by relocating tested seed sources within the species’ range if the targeted site condition matches the adapted climate based on the modeled projections (Gray et al., 2011) and from common garden experiment series (Ding and Brouard, 2022).

Tree improvement and reforestation require accurate estimates of genetic parameters and an understanding of the genetic architecture of growth and adaptive traits. In *P. tremuloides*, estimates of heritability and genetic correlations of adaptive traits are lacking, especially for phenology and stress tolerance, which are important in tree breeding programs to avoid maladaptation and climate-induced mortality in plantations (Ding et al., 2020; Ibanez et al., 2010a; Rohde et al., 2011a). Hänninen (2006) pointed out that unseasonal frosts in spring and fall damage buds and leaves and eventually jeopardize productivity. The genetics of phenology traits are important for tree improvement and reforestation so that the plantation stocks are resilient to climatic-related risks. Numerous breakthroughs in ecological genetics and quantitative genetics are paving the way for successful commercial planting and reforestation with aspen species. Schreiber et al. (2013) used ecological genetics based on among-population genetic variation and demonstrated that assisted migration has high potential to improve reforestation success. Another regional quantitative genetic study showed that sufficient within-population genetic variation leads to adaptation of plantation stocks to the lengthening of the growing season at the juvenile stage without risking severe frost damage (Ding et al., 2020).

Traditional tree improvement can enhance aspen forest adaptation and productivity; moreover, advances in genomics along with groundbreaking work on numerous traits, including phenological traits, stress tolerance (e.g., frost, salinity, drought, heat, UV radiation, etc.), and growth traits, are currently being reported. Because *P. tremuloides* is not yet the true “model species” in genomics research compared to *P. trichocarpa* (Evans et al., 2014; Tuskan et al., 2006), we reviewed some representative studies in *Populus* spp. including sister species that will inspire future studies of *P. tremuloides*. In this review, we carried out a meta-analysis of research papers on adaptive traits with a focus on *P. tremuloides*. Our results cover a broad spectrum of topics in quantitative genetics, ecological genetics, and genomics from 2007 to 2022. This study provides a systematic overview of research on adaptive traits, including studies on mechanisms and genetic variation, as well as molecular regulations and functions related to tree improvement and reforestation. Our findings support key hypotheses and concerns related to adaptive and stress tolerance mechanisms to advance the ‘suite’ of trait improvement strategies under future climatic uncertainties and extremes.

## 2. Adaptive traits in forest trees under climate change

The adaptive capacity of trees is the ability to survive, grow, and reproduce in their natural or reforested habitats and the extent to which the species, population, or family can survive and adapt to a particular sequence of ecological conditions (Dobzhansky, 1956; Mirkena et al., 2010). Adaptive traits are important for domestication, survival, and production involving species in natural, agricultural, and forestry settings (Mirkena et al., 2010). Spring phenological traits are typically assessed adaptive traits in trees, often measured by the timing of bud break or leaf unfolding (Vitasse et al., 2009). Other adaptive characteristics of trees include, but are not limited to, the timing of loss of cold hardiness (Aitken and Adams, 1997) or the timing of dormancy release (Beck et al., 1995). The release of dormancy results in increased respiration rate and photosynthetic efficiency (Beck et al., 1995). The timing of bud set or leaf senescence is another key adaptive trait that marks the onset of growth cessation (Rohde et al., 2011b; St Clair et al., 2005). Leaf senescence can be measured by leaf coloration and leaf abscission (leaf drop) (Fracheboud et al., 2009; Ibanez et al., 2010b). The timing of the onset of frost hardiness is also a critical component of fall phenology (Aitken and Adams, 1997; O’Neill et al., 2001). In addition to these phenological traits, which are relatively easy to evaluate, many other physiological and anatomical traits have adaptive values in terms of drought resistance, salt tolerance, resistance to pest and pathogen, defenses against herbivore, etc. Adaptive traits enable trees to grow and express clines, to grow and reproduce in natural sites and plantations. Developing plantation stocks with adaptive traits matching the site environmental and ecological conditions will ensure future forest performance with long-term economic returns and ecosystem services, as well as improved sustainability values.

Figure 1 shows a schematic model of genetic studies on growth and adaptive traits. Research results on phenotypic variation will guide tree improvement and conservation as well as other future genetic and genomic research by selecting and breeding families and populations with adapted and productive phenotypic performances (e.g., site index, biomass, basal area, and volume) for reforestation. Genetic variation demonstrates the genetic variation within species of multiple populations (e.g., ecotypes, clines, genetic variation among seed zones or breeding regions, etc.); within-populations variation is due to the family variations within seed zones or breeding regions; the among-family variation is usually expressed in the same breeding group, while the genetic variation among individuals or clones is observed in common garden experiments and block trials (Fig. 1). Genotype x environment (GxE) interaction as a quantitative genetic parameter indicates that the same families and clones show different phenotypic rankings at the experimental or plantation sites when the trees are deployed in multiple environments.

Environmental changes are due to both biotic and abiotic stresses, and these are the environmental sources of phenotypic variation. Different environmental stresses cause corresponding or combined variations of phenotypic adaptations. This review focuses on spring and fall phenology, frost hardiness, salinity tolerance, and drought tolerance traits. We will discuss genetic, environmental, and phenotypic variations, including geographical (among population) and within-population variations. Advances in genomics such as marker assisted selection (MAS), genomic selection (GS), genome-wide association studies (GWAS) allow deeper exploration and better utilization of the genetic variation to shorten the long breeding cycles of trees. Genetic analysis of natural populations and breeding populations (in terms of G and GxE) provides insights into molecular regulation and gene functions or interactions with environmental factors to realize genomic applications for multiple trait improvement. Phenotype-genotype associations and genotype-environment (GxE) relationships will combine DNA sequence features with data presented in the three boxes of Fig. 1 to investigate landscape genomics, population structures, and environmental factors to find genomic markers for individual, ecological, and population adaptations and growth dynamics.

**Fig. 1.**
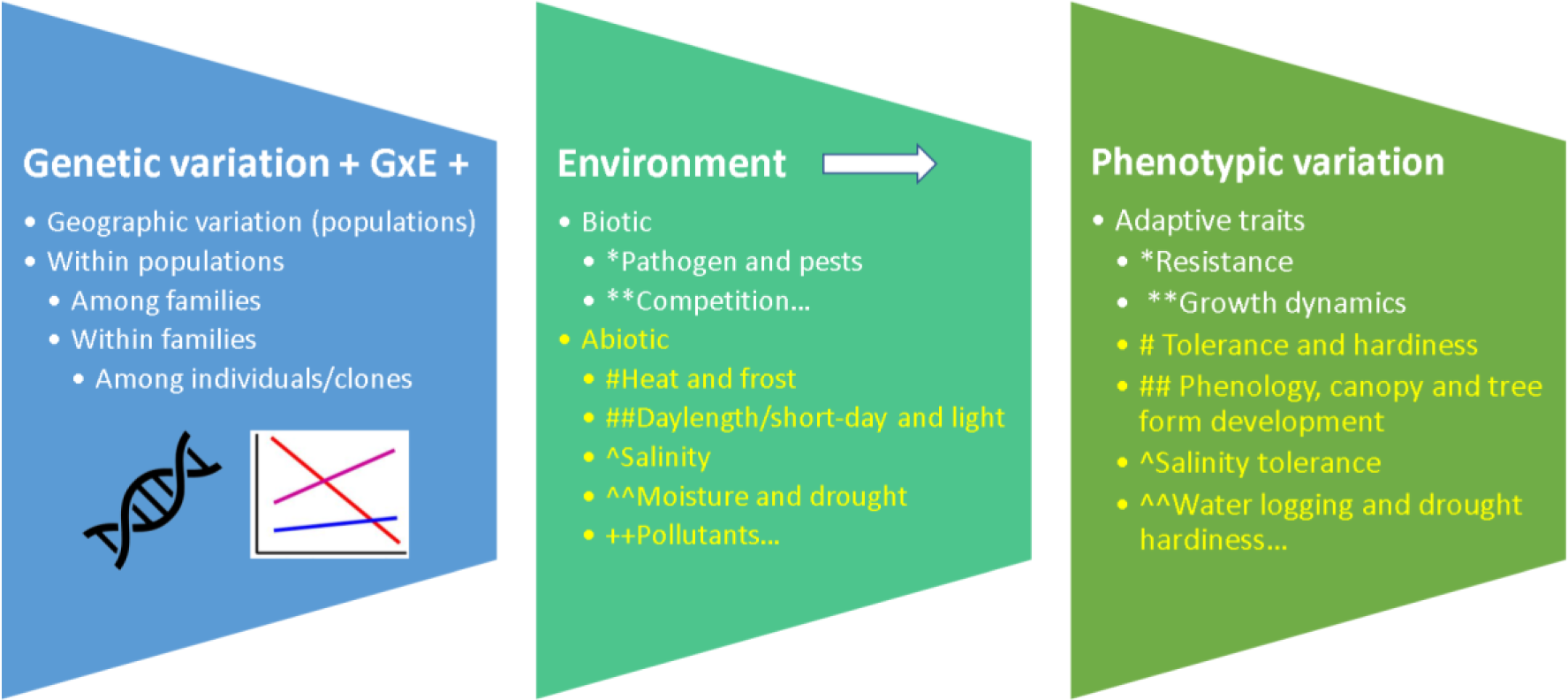
Schematic model for genetic variation in tree growth and adaptive traits. The three boxes present the genetic (G plus GxE), environmental conditions and phenotypic variation (i.e., response) of individual trees.

## 3. Methods

### --Evaluating Geographic, clinal, phenotypic, and genetic variations of adaptive traits

Widespread tree species are usually composed of many locally adapted sub-populations, which are described by a variety of terms, including clines, ecotypes, varieties, or subspecies. A cline is a continuous variation of a species’ genotypes along an environmental gradient and typically refers to observations of a single trait. In widespread tree species, this is a common type of genetic differentiation caused by gene flow patterns. Geographical clines in *Populus* spp. have been reported (McKown et al. 2014), although evidence in *Populus tremuloides* was less frequently found. Clinal variation of the timing of bud break was studied with provenance common gardens previously; for *P. tremuloides*, the clinal variation of spring phenology studied with provenance trials (Li et al., 2010) yielded slopes across western Canada with approximately -0.5 days per degree latitude. Soolanayakanahally et al. (2013) reported among *P. balsamifera*, spanning a latitudinal range from 45 °N to 70°N, negative correlations among bud flush and latitude (*r* = -0.7 to -0.5), equivalent to 1 day earlier per degree of latitude, for a total of 15 days difference between opposite ends of the cline. Allopatric studies of clinal variations within *P. balsamifera* and *P. tremula* demonstrated latitudinal variation in eco-physiological traits such as photosynthesis rate, stomatal conductance, chlorophyll content, and foliar nitrogen (Soolanayakanahally et al., 2015). Geographic clines were also reported for *Phyllocnistis* and *Melampsora* caused parasitic damage of *P. tremula* in Sweden (Albrectsen et al., 2010).

### 3.1 Spring phenology

Spring bud break of trees is the initiation of shoot growth at the apical meristem and the formation of new leaves after the release from winter dormancy (van Volkenhrugh and Taylor, 1996). The timing of bud break in spring is usually controlled by two major mechanisms: a chilling requirement and a heat sum requirement (Korner and Basler, 2010). Temperate trees have moderate to strong chilling requirement before heat accumulation begins (Polgar et al., 2014). Boreal trees such as *P. tremuloides* break bud mainly in response to accumulated heat (Li et al., 2010). In *P. trichocarpa*, the timing of bud break and leaf flush differed by approximately 16-18 days among the most northern and southern genotypes (44° N to 60° N latitude) with 1-2 days earlier per degree latitude to the north, although the relationship could not be proven with R–squared values (McKown et al., 2013). However, no trends in bud flush were found in the Swedish aspen collection along the transect from 56° N to the 66° N latitude (Luquez et al., 2008). Bud-break mechanisms will inform breeding and conservation decisions; the molecular regulator and pathways related to the poplar *EARLY BUD-BREAK 1* (*EBB1*) gene, which is a positive regulator of bud-break and encodes a transcription factor from the AP2/ERF family (Busov et al., 2016; Yordanov et al., 2014), are frequently studied. Quantitative trait loci (QTLs) information of *P. trichocarpa* for adaptive traits including spring and fall phenology are published (http://www.phytozome.net/poplar) and benefits not only the comparative genomics studies of *Populus* species but also *Salix* species (Ghelardini et al., 2014).

### 3.2 Fall phenology

In autumn phenology, the timing of bud set is a complex physiological process and trees respond to shortened daylight by regulating endogenous phytohormones such as cytokinins and gibberellic acid (Horvath et al., 2003). In *P. tremula*, genetic variation in phytochrome genes has been associated with latitudinal clines in bud set (Ingvarsson et al., 2006). Genes related to abscisic acid production have been shown to play roles in regulating bud set in poplar trees (Rohde et al., 2002). Bud set timing is often correlated with growth, representing a tradeoff between utilizing the growing season for photosynthesis and timely recycling of nutrients before fall frosts (Bohlenius et al., 2006). Leaf senescence is not as affected by photoperiod as bud set (Michelson et al., 2018), although both leaf senescence and bud set are treated as autumn phenology here. The fall leaf color and leaf fall are easier to observe than terminal bud set. Multiple pigments, specifically anthocyanins, carotenoids, and chlorophyll, cause leaf coloration. Carotenoids degrade while anthocyanin accumulates during fall color (Keskitalo et al., 2005). Chlorophyll drastically decreases in plastids and vacuoles of mesophylls until the cytoplasm is gone (Keskitalo et al., 2005). In *Populus* trees, autumn coloring often shows yellow and occasionally red hues that appear before leaf fall. However, species such as *Ulmus americana* only express yellow and brown colors. High night temperatures may delay fall coloration under the same photoperiod. The correct timing of leaf coloring and leaf shedding is an important mechanism for the recycling of plant resources and nutrients (Tanino et al., 2010). Leaf protein relocation happens during abscission, which helps plants to overwinter as nitrogen moves into bark reserves (Cooke and Weih, 2005), while carbon nutrients are lost when the leaf falls. Thus, this trade-off between seasonal nitrogen cycling and carbohydrate production is likely to occur, and hence the optimal timing of leaf senescence is important for acclimation and ultimately adaptation in fall.

There is usually a strong clinal variation in autumn phenology. Ingvarsson et al. (2006) reported that bud set of *Populus tremula* in Sweden over a latitude of 10° was 4.4 days earlier per latitude further north (R² = 0.90). Similar studies in Sweden showed about 3-4 days earlier bud set per latitude northwards with R² = 0.63∼0.68 (Luquez et al., 2008). In Canada and Alaska, Soolanayakanahally (2013) found that the date of bud initiation was 4 days earlier per degree latitude further north, with R² values of 0.60-0.90. Fracheboud et al. (2009) found that the onset of leaf senescence in northern *P. tremula* provenances from 56°N to 66°N was ∼3.9 days per degree latitude earlier after August 29 (R^2^=0.45, *P*<0.001) and that bud set had a higher latitudinal correlation (R^2^=0.82, *P*=0.001) than leaf senescence. The timing of leaf coloration is highly correlated with latitude. Soolanayakanahally (2013) found a two-day lag per degree latitude for leaf coloration phenology from northern to eastern provenances of *Populus balsamifera*, for which leaf coloration lasted for 1.5-2 months.

The physiological mechanisms that control the timing of leaf coloration and leaf senescence involve phytochromes, a group of pigments that sense differences in light quality and night length. For example, two forms of Phytochrome A vary between P_r_ and P_fr_ when exposed to red or far-red radiation (Sharrock, 2008). Upon exposure to red light, P_r_ converts to P_fr_ to regulate gene expression and alter plant morphological traits, such as stem, hypocotyl or shoot elongation, seed germination, flowering, etc. In temperate plants, the P_fr_/P_r_ ratio is used as a measure of daylength because P_fr_ is converted back to P_r_ in darkness. This indicator is sometimes combined with additional temperature and moisture cues to initiate fall phenological traits such as bud set, leaf abscission, and dormancy (Sharrock, 2008). Short daylength and low temperature are different cues for cold acclimation, and the *PHYTOCHROME A* gene is important for photoperiodic regulation of abscisic acid and dehydrins (cue for short daylength), which are critical for activating acclimation in hybrid aspen (Welling et al., 2002). Figures 2-3 below illustrate the methodology for measuring spring and fall phenology as well as frost hardiness in fall for *P. tremuloides* (Ding, 2015).

**Fig. 2.**
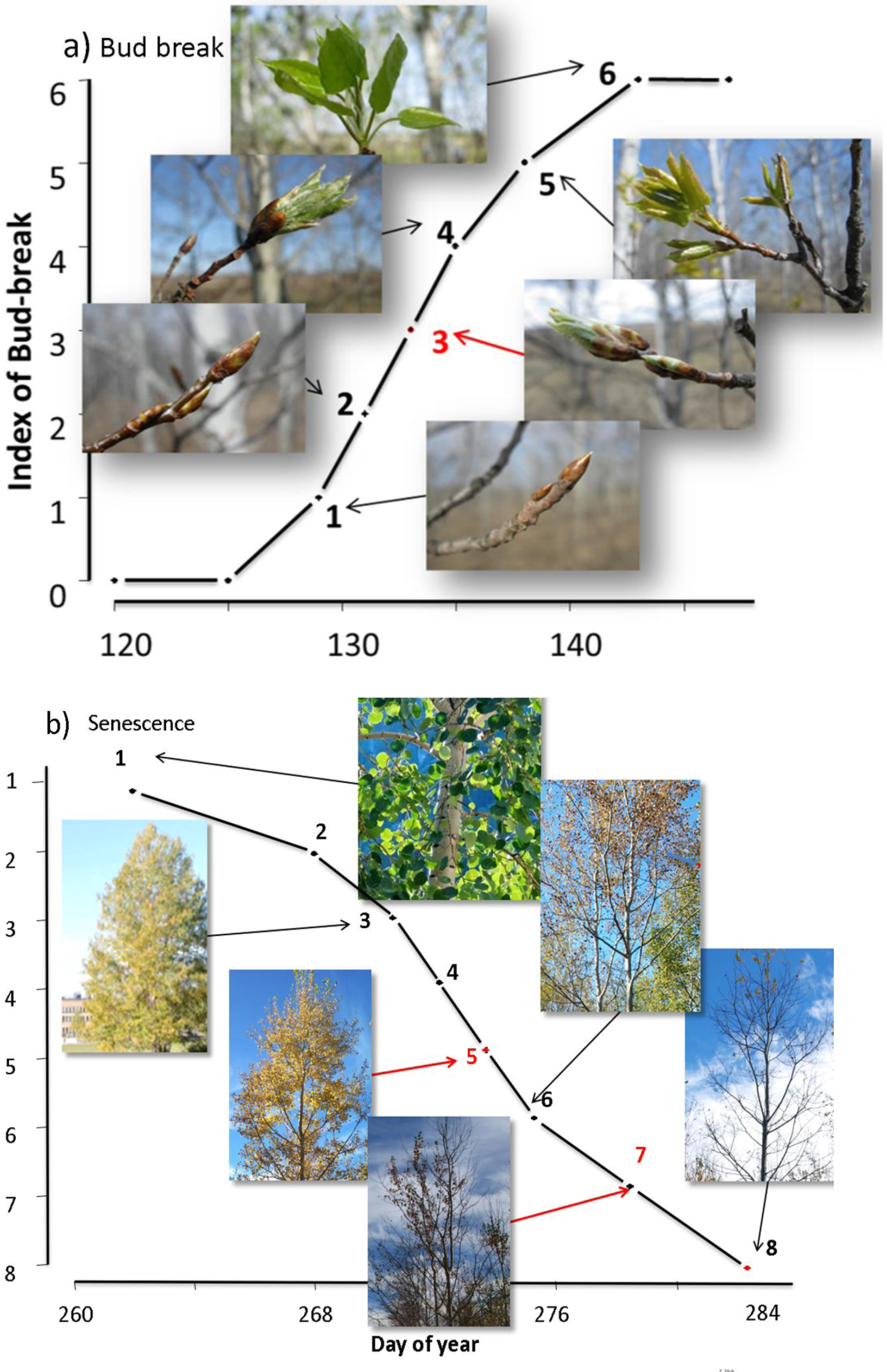
Scoring indices of spring bud break and fall leaf senescence in *Populus tremuloides*. Notes: The x-axis is the Day-of-Year and the y-axis is the index of score. Both methods are semi-categorical. Scoring schemes are for a) spring phenology and b) fall phenology based on the studies by Ding (2015). The typical phenology measurement is based on visual observation which can be replaced with remote sensing methods especially in the controlled environment of greenhouses. The phenology stages of 5 and 7 can be used as critical scores to determine field phenology.

**Fig. 3.**
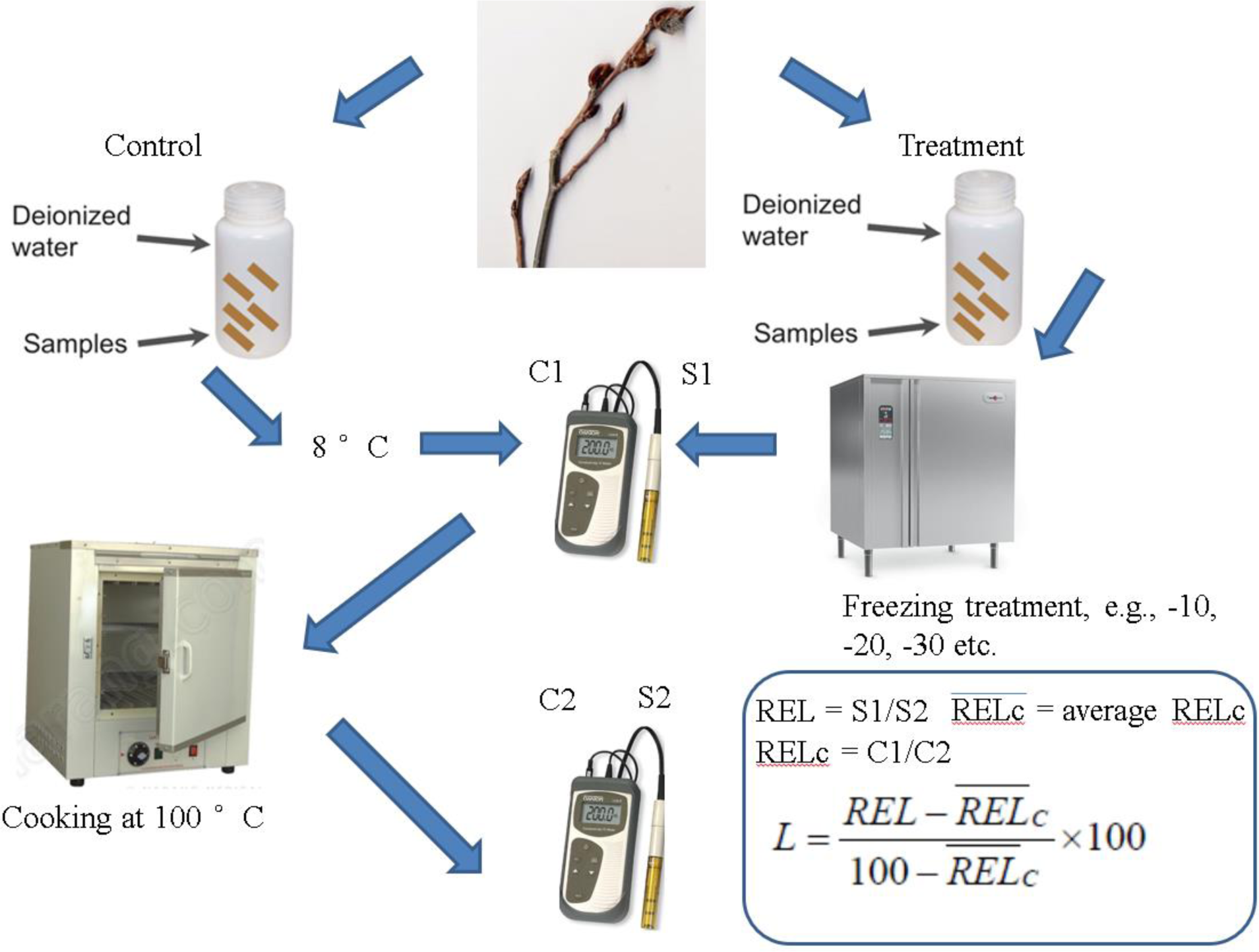
The artificial freezing test method for measuring frost hardiness in *P. tremuloides*. Notes: C1, conductivity reading *before* samples are boiled/cooked at 100°C for 50 mins; C2, conductivity reading *after* samples are cooked at 100°C; REL, the relative amount of electrolyte leakage of samples subjected to the treatments, calculated as (c1/c2)*100; L is the ratio of cell lysis (%); *R̅E̅L̅*_*C*_ the average relative amount of electrolyte leakage of the controls. S1 is the conductivity reading after the freezing treatment under different temperatures; S2 is the conductivity value after the samples were cooked after the freezing treatment.

### 3.3 Frost hardiness and clinal variation

Frost hardiness traits also have strong geographical clines, and the onset of cold hardiness is typically correlated with bud phenology or leaf senescence (Howe et al., 2003). The critical temperatures for 50% lethal damage (LT50) of trees under an artificial freezing test are used to distinguish the hardiness variation among populations or provenances (Larcher, 2003), although sometimes 50% damage may not be achieved in all experiments. Schreiber et al. (2013) found that the LT50 of *P. tremuloides* reached below -20°C to -50 °C from August to late September and strong deviations were found between the British Columbian (northern) and Minnesota (southern) provenances. There are many studies of hardiness and onset of hardiness in trees other than *P. tremuloides*. In *Alnus sinuata*, Benowicz et al. (1999) found that LT50 in January reached a range of -50°C to -20°C showing low difference among populations for fall frost injury. Aitken and Adams (1996) demonstrated that in coastal and Cascades populations of *Pseudotsuga menziesii*, frost hardiness in November of the needles and buds was higher for Cascade populations, which were about 3∼20% less vulnerable to frost than the coastal populations. However, the population divergence of tissue damage was small in January (0∼ 9% for buds, stems and needles). For *Tsuga heterophylla*, Oregon populations showed higher fall frost injury (44∼52%) and spring frost damage of needles (45∼48%) than British Columbian populations (north) for which the injury was 21∼29% in fall and 28∼40% in spring, respectively, under a freezing temperature from -6 °C to -14 °C (Hannerz et al., 1999). In *Pinus sylvestris*, the maximum hardiness of needles varies from population to population: for instance, damaging temperatures reached down to -40°C in the northern provenance, while all populations in Finland were frost hardy at -30°C (Hurme et al., 1997). For altitudinal clines, Rehfeldt (1988) summarized provenance test results of 173 *Pinus contorta* populations in the Rocky Mountains from 1,300 m to 2,900 m and found a 0.0009 decrease of the frost injury index per meter altitude when populations are from higher origins. For *Picea sichensis*, another coastal conifer, Mimura and Aitken (2007) reported strong latitudinal clines of cold injury index in a range-wide collection area along the Pacific coast from Alaska to British Columbia, with 7% higher hardiness per 100 km from south to north (R^2^=0.58, *P*<0.0001).

The application of genomics and field common garden experiments demonstrated finer spatial and temporal population structures and divergence in frost hardiness and other adaptive traits. Zhang et al. (2019) investigated 451 *P. trichocarpa* genotypes with molecular genetics means and in field common gardens and suggested that the cold injury and the bud set timing (DoY) significantly decreased when the latitude increases (R=0.18, P<0.001); they also identified 172 candidate genes for cold hardiness and found that *Potri.017G033100* (GPX3; glutathione peroxidase 3) was associated with timing of bud flush and cold hardiness along the latitudinal gradient. Landscape genomics and genomic methods (e.g., redundancy analysis) will support similar studies in aspen, *Populus,* and other forest tree species (Capblancq and Forester, 2021; Wang et al., 2022). Other tools also can be used in adaptive trait studies including epigenetics tools and microRNAs (Ma et al., 2018; Zhang et al., 2022), although such studies are still not widely reported in aspen trees.

### 3.4 Salinity stress

Salinity stress is a major abiotic pressure that challenges reforestation. Approximately 99% of flora in the world cannot survive in high-salt environments which total to 8.3 billion ha globally (Flowers and Colmer, 2008; Zhang et al., 2021; Zhu, 2001). Salt stress leads to reductions in growth, productivity, and survival in numerous plant species (Munns and Tester, 2008). The stress induced by salinity is caused by three mechanisms: osmotic stress due to a more negative soil water potential, accumulation of toxic ions, and disturbances in nutrient balance (Munns and Tester, 2008).

Halophytic plants are salt-tolerant populations, families, or varieties that can tolerate and reproduce in mid to high salinity soils (sodium chloride > 200mM). However, most of the woody plants belong to glycophytes whose growth is inhibited by NaCl concentrations, even when exposed for a short period of time (Acosta-Motos et al., 2017; Cheeseman, 2015). It has been reported that concentrations as low as 20-mM NaCl can inhibit growth in many glycophytes (Flowers, 2004; Renault et al., 2001). Traditional methods for the improvement of tree afforestation success in salt-affected areas involve various soil improvement strategies, e.g., selecting clones well adapted to soil salinity or changing the soil conditions. Nonetheless, the most cost-effective approach is screening and breeding salt-tolerant plants with genomics, quantitative trait loci (QTL) mapping, genomic prediction (GP), and genome editing (GE), and those approaches have been used to accelerate traditional poplar breeding (Biselli et al., 2022). The transgenic approach for improving salinity tolerance has been widely used in crop plants. In *Vigna aconitifolia* and *Panicum virgatum*, the transformed pyrroline-5-carboxylate synthetase (*P5CS*) gene encoding for the rate limiting enzyme important for proline biosynthesis enhanced salt tolerance (Guan et al., 2018; Surekha et al., 2014). To enhance the salt tolerance of *Eucalyptus globulus* trees, a *12oda* gene encoding for choline oxidase was introduced and improved salt tolerance (Matsunaga et al., 2012). Current studies focus on gene editing to enhance *Populus* salt-tolerance. In a hybrid poplar study, the H^+^ - pyrophosphatase gene (*PtVP1.1*) of *Populus trichocarpa* improved salinity tolerance in *P. davidiana* × *P. bolleana* plants after *PtVP1.1* overexpression (Yang et al., 2015). The *LbDREB*-transgenic *Populus ussuriensis* (incorporating a *DREB* gene from the halophyte *Limonium bicolor*) exhibited enhanced salt resistance by increasing its reactive oxygen species (ROS), reducing its malondialdehyde (MDA) content, and increasing its proline accumulation in leaves (Zhao et al., 2018). The overexpression of *PtPCBER* promoted salt tolerance in transgenic poplar by increasing ROS scavenging capacity (Wei et al., 2022). Therefore, the transcription factors and molecular mechanisms underlying salt tolerance are an important key to the enhancement of salt resistance in *Populus*.

### 3.5 Drought tolerance

Drought stress is the leading environmental factor that impacts growth and productivity of plants; water availability is among the most limiting factors to plant growth worldwide (Almeida-Rodriguez et al., 2010). Drought is classified into three types (Dai, 2011; Wilhite, 2000). The most common type of drought is the *meteorological drought*, which is a prolonged period with low precipitation. Meteorological drought is often accompanied by above-normal temperatures and precedes and causes other types of droughts. An *agricultural drought* is a period with dry soils (lower soil water potential) that results from below-average precipitation, intense but less frequent rain events, or above-normal evaporation, all of which lead to reduced crop production and plant growth. A *hydrological drought* occurs when river streamflow and water storages in aquifers, lakes, or reservoirs fall below long-term mean levels; such drought develops more slowly, whereby stored water is depleted but not replenished.

Plants resist drought stress through various morphological, biochemical, and physiological mechanisms (Farooq et al., 2009). Morphologically, plants resist water deficit mainly through drought escape, drought avoidance and drought tolerance. *Drought escape* is accomplished through a shortened life cycle or growing season, allowing plants to complete their life cycle before the environment becomes dry. Drought escape occurs when the plants’ phenological development periods are completed before terminal drought stress predominates (Araus et al., 2002). *Drought avoidance* is the ability of plants to avoid reduced tissue water content despite reduced water content in the soils (Basu et al., 2016). Under transient periods of drought stress, drought avoidance occurs when plants increase water-use efficiency, lower stomatal conductance, reduce leaf area, limit vegetative growth, change hydraulic conductivity, or increase root growth to avoid dehydration (Kooyers, 2015). Isohydric (drought-avoidant) plants control water loss mainly through control of stomatal movements (Almeida-Rodriguez et al., 2010). Common strategies include reduction of transpiration area through leaf shedding and production of smaller leaves to reduce the water loss, deposition of heavy cuticle layer on the epidermis, sunken stomata, and trichomes (Farooq et al., 2009). Root characteristics such as increased length, density and depth are the main drought avoidance traits that contribute to plant growth under drought stress (Turner et al., 2001).

*Drought tolerance* involves enduring low tissue water content through adaptive traits by maintaining cell turgor through osmotic adjustment and cellular elasticity as well as increasing protoplasmic resistance (Morgan, 1984). Physiologically, osmotic adjustment, osmoprotection, antioxidation, and the scavenging defense system are the most important bases for drought tolerance (Farooq et al., 2012). Under drought stress conditions, sugars and free amino acids have been reported to accumulate in many plant species (Manivannan et al., 2007; Sankar et al., 2007). Besides lowering water potential and increasing cell turgor by osmotic adjustment, compatible solutes can also protect enzymes from the damaging effects of Reactive Oxygen Species (ROS) (Farooq et al., 2012). Similar to salt stress described above, in response to ROS caused by drought stress, complex antioxidant enzyme systems such as superoxide dismutase (SOD), peroxidase (POD), catalase (CAT) and ascorbate peroxidase (APX) are triggered in plants (Kaya et al., 2006; Vardharajula et al., 2011). In addition to the high activity of antioxidant enzymes, some non-enzyme antioxidants such as glutathione (GSH), ascorbate (ASC) and carotenoids also provide protection by quenching toxic ROS in plants (Kubiś et al., 2014; Singh et al., 2015). The antioxidant enzymes may directly scavenge ROS or may produce non-enzymatic antioxidants, which can act to protect the integrity of the photosynthetic membranes under oxidative stress (Shakeel et al., 2011). There are many drought tolerance discoveries in plant species or crop sciences other than in *Populus* studies, e.g., the drought-tolerant variety of sweet sorghum (*Sorghum bicolor*) had significantly higher activities of the antioxidant enzymes than the drought-sensitive variety (Guo et al., 2018). Also, drought stress of soybean (*Glycine max*) triggered higher activities of CAT, POD, APX and glutathione reductase enzymes in the drought-tolerant variety compared with the drought-sensitive variety (Devi and Giridhar, 2015). Similar results were also found in faba bean (*Vicia faba*), with the activities of antioxidant enzymes being higher in the drought-tolerant genotype compared with the drought-sensitive plants (Abid et al., 2017). Therefore, the ability of antioxidant enzymes to scavenge ROS is among the most important mechanisms of drought tolerance in plants that have not yet been studied in detail in *Populus* trees.

Genomic and genetic variations of drought tolerance were explored in multiple *Populus* species, although less work on aspens has been done to date. Chamaillard et al. (2011) examined 30 genotypes of *P. nigra* and found that the drought effect was neither genotype-nor population-dependent and that showed the same ranking of populations maintained at multiple sites. Liu and El-Kassaby (2019) investigated the plasticity of drought tolerance and phenological traits in *P. trichocarpa* and demonstrated that higher drought resistance, extended bud set and growth period as well as a shorter post-bud set period to leaf drop link to greater height gain under a future extreme climate that includes factors such as drought. In a genome-wide association study (GWAS) also conducted in *P. trichocarpa*, two genes out of 20 gene models were reported: *Potri.001G411800* encoding a EF-hand calcium-binding domain containing protein and *Potri.009G158900* encoding a late embryogenesis abundant (LEA) hydroxyproline-rich glycoprotein; these gene were identified to have major roles in responses to drought, salinity, osmotic and temperature-related stresses (Chhetri et al., 2019). In *P. euphratica*, for example, *LAC2* (*LACCASE*) was found to be associated with improved drought tolerance (Niu et al., 2021). Sex-specific responses in drought adaptation have also been studied in *Populus* spp. (Melnikova et al., 2017), and males trees were reported to be more adaptive to environmental stress conditions with less damage, better growth, and higher photosynthetic capacity and antioxidant activity than females.

## 4 Molecular genetics and genomic tools for studying *Populus tremuloides* and *P. tremula*

Advances in forest genomics have created great opportunities for discovery and provided many new molecular-level traits that may be utilized in tree improvement. Some of these new opportunities include: 1) manipulate species’ genome architecture and components for increasing productivity, adaptation, and resilience; 2) use marker-assisted selection and molecular breeding by exploring and employing markers and candidate genes such as those underlying the QTLs (Ghelardini et al., 2014); 3) exploit genome-wide associations (GWAS) and genomic selection to accelerate trait and population improvement (Borthakur et al., 2022; Ding et al., 2018); and, 4) utilize gene discovery, epigenetics, and functional studies for gene editing, complex trait design etc. (Borthakur et al., 2022). Although the genomes of *P. trichocarpa* and eucalyptus have been studied and used far more in molecular studies of trees, aspen has a good chance of being a future “model” - as the tool of genomic studies becomes more feasible - because of the strong genetic variation in aspen’s adaptive traits and growth. Previous studies of adaptive traits have focused on spring and fall phenology and on stress tolerance.

Pioneering studies in *P. trichocarpa* uncovered the genome-wide association between single nucleotide polymorphism (SNP) markers and local phenological and eco-physiological adaptation traits as well as growth traits. Among 29,355 filtered SNPs representing 3,518 genes, 240 genes were associated with phenology (McKown et al., 2014). Bud-break was associated with 16 genes and southern- and northern populations broke bud earlier than those from the center (McKown et al., 2018). Environmental variables were employed with GWAS and half of the adaptive phenology and eco-physiology traits were significant, indicating population adaptive variation based on *Q*_ST_; selection signals were shown based on 2,855 SNPs, and among which 118 SNPs within 81 genes were also associated with adaptive traits such as autumn phenology, height, and disease resistance (Porth et al., 2015). In *P. deltoides*, subpopulation differentiation was low (*F*_ST_ = 0.022-0.106) while genetic diversity was high with a large effective population size (Ne ∼32,900), and levels of linkage disequilibrium low to moderate (Fahrenkrog et al., 2017).

Those findings are novel and applicable in landscape genomics and phylogenomic studies of aspen (Balkenhol et al., 2019). Little genetic structure was found in *P. tremuloides* based on microsatellite markers (Latutrie et al., 2016). With a more advanced sampling range and genomic analyses, four main genetic lineages (clusters) were demonstrated for this North American species’ range and frequent triploids were identified in western USA, Mexico and western Canada, in regions more prone to drought (Goessen et al., 2022).

## 5 Genetic engineering strategies for developing climate-smart quaking aspen

Climate change aggravates the adverse effects of biotic stresses (fungal, bacterial, and pest, etc.) and abiotic stresses (drought, cold, heat, etc.) (Ahuja et al., 2010; Zandalinas et al., 2021). Poplar trees are even more vulnerable to the climate-induced recurrence of these multiple biotic and abiotic stress because of their several decades long life cycle (Sturrock et al., 2011; Naidoo et al., 2019; Biselli et al., 2022). Genetic engineering is one of the most potent and promising techniques to create climate-resilient poplar clones used in plantations (Hu et al., 2014; Thakur et al., 2021; Zhang et al., 2022). For example, in China, the genetically engineered poplar clones with Bt expression resulted in a substantial 90 % reduction in leaf damage suffered from pests compared to unmodified wild-type clones (Wang et al., 1996) and have been commercialized and planted to occupy 450 hectares until 2011 (Lu and Hu, 2011; Wang et al., 2018). To deal with the aggravated stresses of poplars from climate change, many genetically engineered elite lines have been generated in many poplar species, such as *P. tremuloide*s and its hybrids (reviewed in Naidoo et al., 2019; Thakur, Ajay K., et al., 2021; Borthakur et al., 2022).

With climate change, weed pressure poses more threats to forest production due to weed infestation, which imposes a competition for nutrient sources with poplar in forestry systems (Otto et al., 2010; Watt et al., 2019; Fuchylo et al.,2022). Therefore, developing a strategy that reduces the harmful effects of weeds without affecting poplar is needed. Through the genetic engineering approach, transgenic poplars expressing herbicide resistance genes are generated, which enables these poplars to remain unaffected in forests with the application of weedicides (herbicides). Multiple *P. tremula* × *P. tremuloides* transgenic lines have been generated expressing the herbicide-resistant proteins ENOYLPYRUVYLSHIKIMATE-3-PHOSPHATE SYNTHASE (*EPSPS*) from Agrobacterium or glyphosate oxidoreductase (*GOX*), for conferring tolerance to the Roundup® herbicide whose active ingredient is glyphosate (Melian et al., 2002). Hybrid *Populus* (*P. tremula* × *P. tremuloides*) expressing *CP4 EPSPS* for resistance against glyphosate suffered little damage even after being subject to the treatment of 3.9 kg of glyphosate per hectare, while the growth of the weeds is inhibited in the same herbicide treatment (Melian et al., 2002). Together with the promising results from the overexpression of the herbicide-resistant genes in other poplar species mitigating the adverse effect of weeds (Fillatti et al., 1987; Review in Dubouzet et al., 2013; Review in Thakur, Ajay K., et al., 2021), genetic engineering with herbicides resistant genes is an efficient approach coping with the weed problem enhanced by climate change.

With extreme weather induced by climate change, poplar trees are increasingly susceptible to various types of abiotic stresses like drought, salinity, heat, and metal toxicity (Seserman et al., 2018; Dănilă et al., 2022). These stresses induce multiple molecular and physiological changes in the tree, resulting in multiple undesirable responses, such as ion toxicity, active oxygen damage and photo-inhibition in plants (Harfouche et al., 2014; Zhu et al., 2016). These stress responses could limit the productivity and health of planted poplars (Marron et al., 2014). To cope with the challenge of these abiotic stresses, multiple genetic engineering approaches have been implemented in poplar species (reviewed in Naidoo et al., 2019; Thakur, Ajay K., et al., 2021; Borthakur et al., 2022). For example, *P. tremula × P. tremuloides* transgenics were created that confer enhanced drought tolerance by overexpressing *Arabidopsis* stress-responsive galactinol synthase (*AtGolS2*) (Shikakura et al., 2022), prevent cold acclimatization by ectopic expression of the oat *phytochrome A* (Olsen et al., 1997), and strengthen salt tolerance by overexpression of *P. trichocarpa* salt overly sensitive 2 (*PtrSOS2*) proteins (Zhou et al., 2014). With the functional characterization and increasingly new knowledge of abiotic stress tolerance-related genes as a resource for genetic engineering in poplar, developing transgenic poplars with the modification of several genes at once for increasing multiple stress tolerance levels could be desirable to enhance poplar productivity without sacrificing its health.

Many types of diseases and pests affecting forests are anticipated to become more widespread with climate change (Desprez-Loustau et al., 2007; Sturrock et al., 2011; Fischer et al., 2019; Linnakoski et al., 2019). For example, severe drought in west-central Canada induced by El Niño (Hogg and Michael, 2015; Yang et al., 2020) resulted in the incidence of decay fungi (*Phellinus tremulae*) whose outbreak could result in aspen heart rot. Overexpression of pinosylvin synthase in transgenic aspen line H4 was shown to enhance its resistance to *Phellinus tremulae* (Seppänen et al., 2004). Although we need more examples showing that the genetically engineered *Populus tremuloides* or its hybrids could cope with more pests and diseases, many such examples are shown in other poplar species already (reviewed in Naidoo et al., 2019; Dort et al., 2020; Thakur, Ajay K., et al., 2021; Borthakur et al., 2022). These results could demonstrate the capability and strength of genetic engineering in fighting biotic stresses affecting poplar species.

With more detailed annotation for an increasing number of poplar genomes (including that of aspen), the fast development of genetic modification tools such as CRISPR/Cas9, and better functional characterization of the poplar genes, the genetic engineering approaches could be integrated into the poplar improvement program to generate the climate-smart poplar elite lines (reviewed in Bewg et al., 2018; Goralogia et al., 2021; Cao et al., 2022). Additionally, given that DNA-free genome edits are not regulated as GMO, transgene-free trees generated by optimizing CRISPR-based technology might speed up the rapid deployment and large-scale field plantation of these genetically “optimized” trees in the U.S.A. (Strauss et al., 2019; Anders et al., 2023).

## 6 Meta-analysis of the genetics of adaptive traits, frost, salinity, and drought tolerance

*Populus* spp. are becoming the model species to explore and use the within-and among population/family/variety variation to resolve the deforestation crisis and improve the sustainability of plantation forestry. *P. tremuloides* was not the major focus in genomics originally; but the genetics of adaptive traits in general, although for sister species in this genus, have been researched. In this section, we therefore extended the species scope to include possible examples for future research in aspen.

We explored previous research done on *Populus* tree adaptation by searching the Web of Science search engine for peer-reviewed literature during 2001-2022 with the key words “*Populus*, adaptive trait, genomics”. There were 65 core literature citations filtered out (Table 1). We categorized the retrieved research items in terms of their adaptive traits and species focus (Figures 4-6). Phenology and genomics studies (e.g., genome research, bioinformatics, gene expression, genetic engineering, etc.) outnumbered the other studies on non-phenological adaptive traits such as tolerance to various stresses including frost, drought, salinity, UV, flood, etc. (Fig. 4). Eco-physiological traits will be important in future research to explain underlying causes related to climatic stress and phenological responses. Approximately 20% of the surveyed studies were done on the two “model” *Populus* species *P. trichocarpa* and *P. deltoides*, including their hybrids. But as can be seen, aspen trees were considerably lagging behind in terms of research focus compared with those species. However, hybrids and pure species of aspens have sparked increasing interest through forest genetics and genomics studies for a decade now, especially at the regional and continental scales (Goessen et al., 2022; Latutrie et al., 2016; Mock et al., 2012). Our literature analysis did not reveal a strong overall trend of increasing findings or publications (R^2^=0.18, Fig. 7). As poplar species studies were not correlated with time (Fig. 6), research interests seemed not to have changed over time. Also, multiple *Populus* species were covered in previous studies other than *P. trichocarpa* and *P. deltoides* and their hybrids. Increasingly, resources become available to promote future aspen studies. For example, facilities such as the Centre of Bioenergy Innovation, Department of Energy USA, are pioneers in *Populus* genomic and phenomics research, including the coastal and inland field test series of common gardens covering thousands of genotypes, genotyping and transcriptomic platform (https://cbi.ornl.gov/). These types of resources are promising to be influential and supportive.

**Fig. 4.**
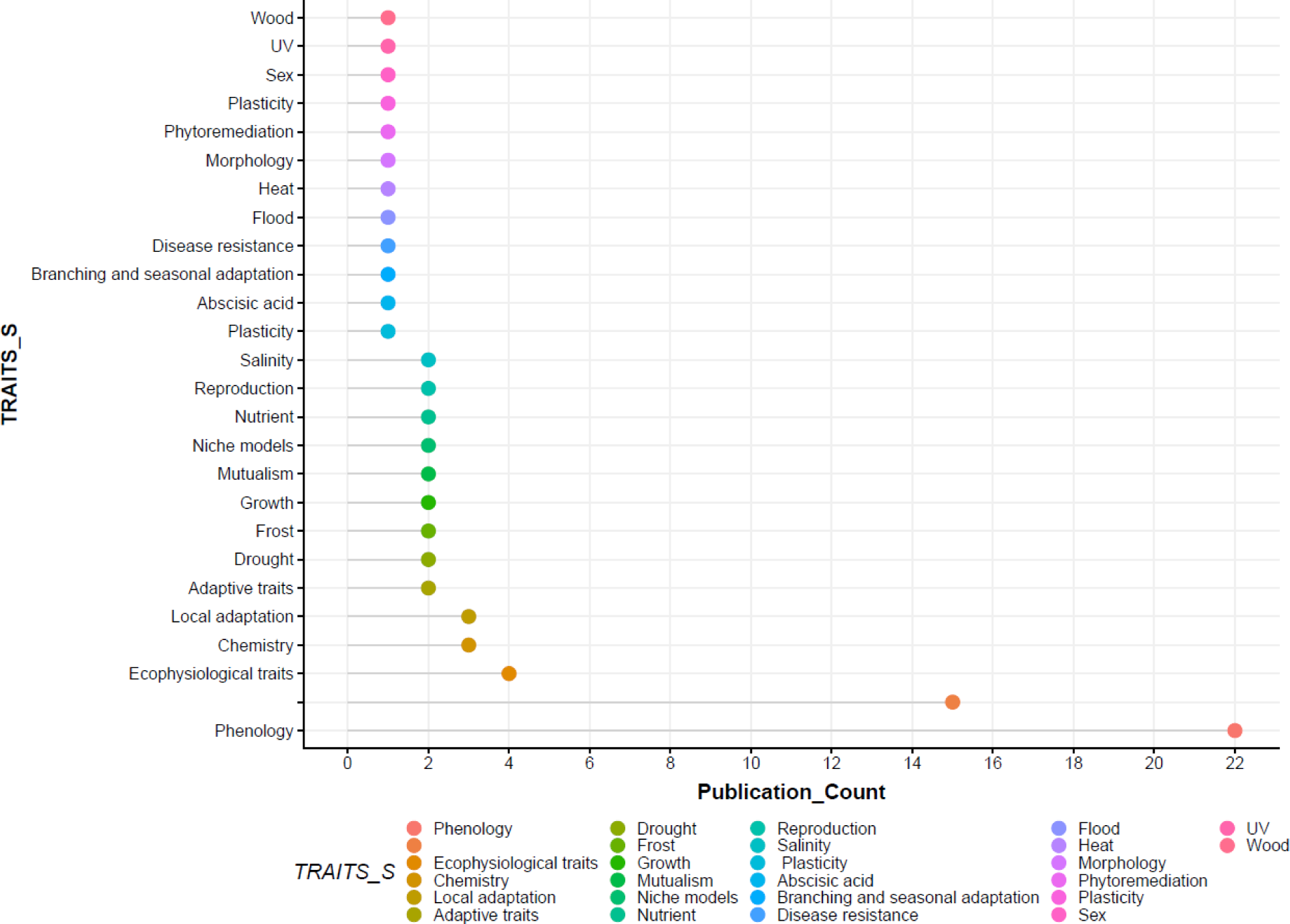
The number of publications in *Populus spp*. from 2001 to 2021 in broad spectra of adaptive traits. Notes, the first dot (no name with color code) indicates not a specific trait study that can be categorized like the others.

**Fig. 5.**
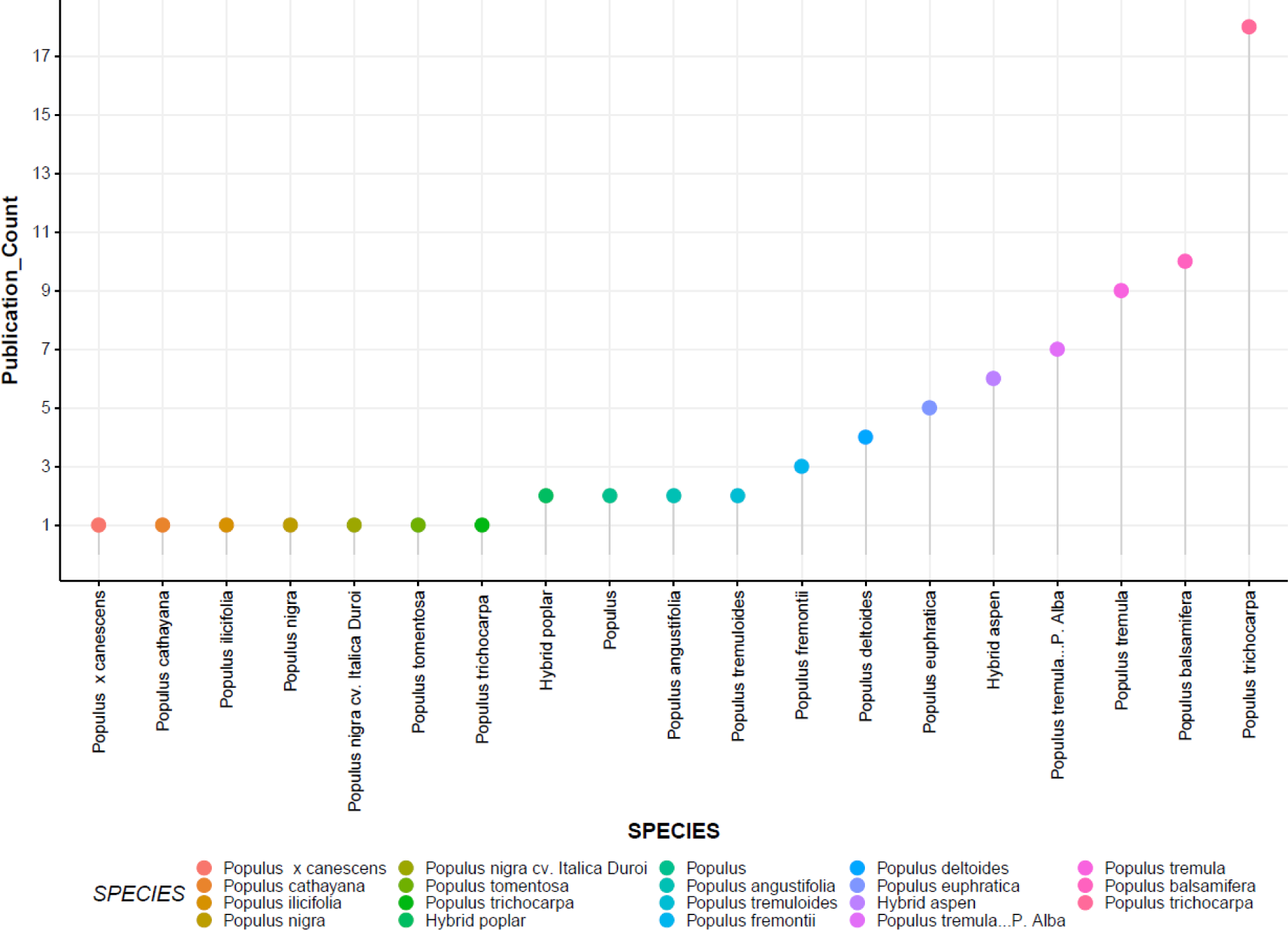
The number of publications in *Populus spp*. from 2001 to 2021 across broad spectra of adaptive traits.

**Fig. 6.**
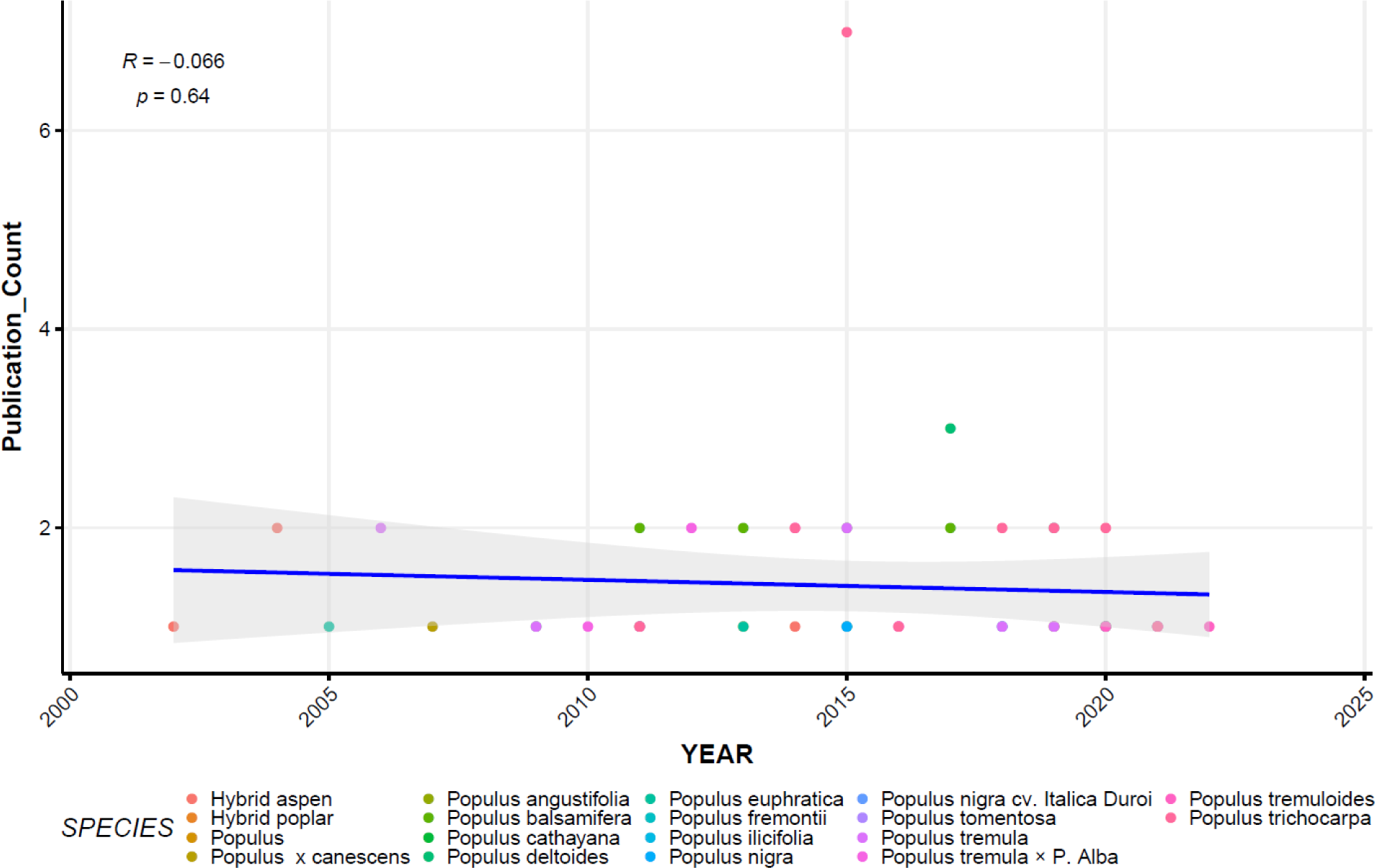
The temporal trend of publications regarding *Populus* spp. and adaptive traits from 2001 to 2021 with a regression line.

**Fig. 7.**
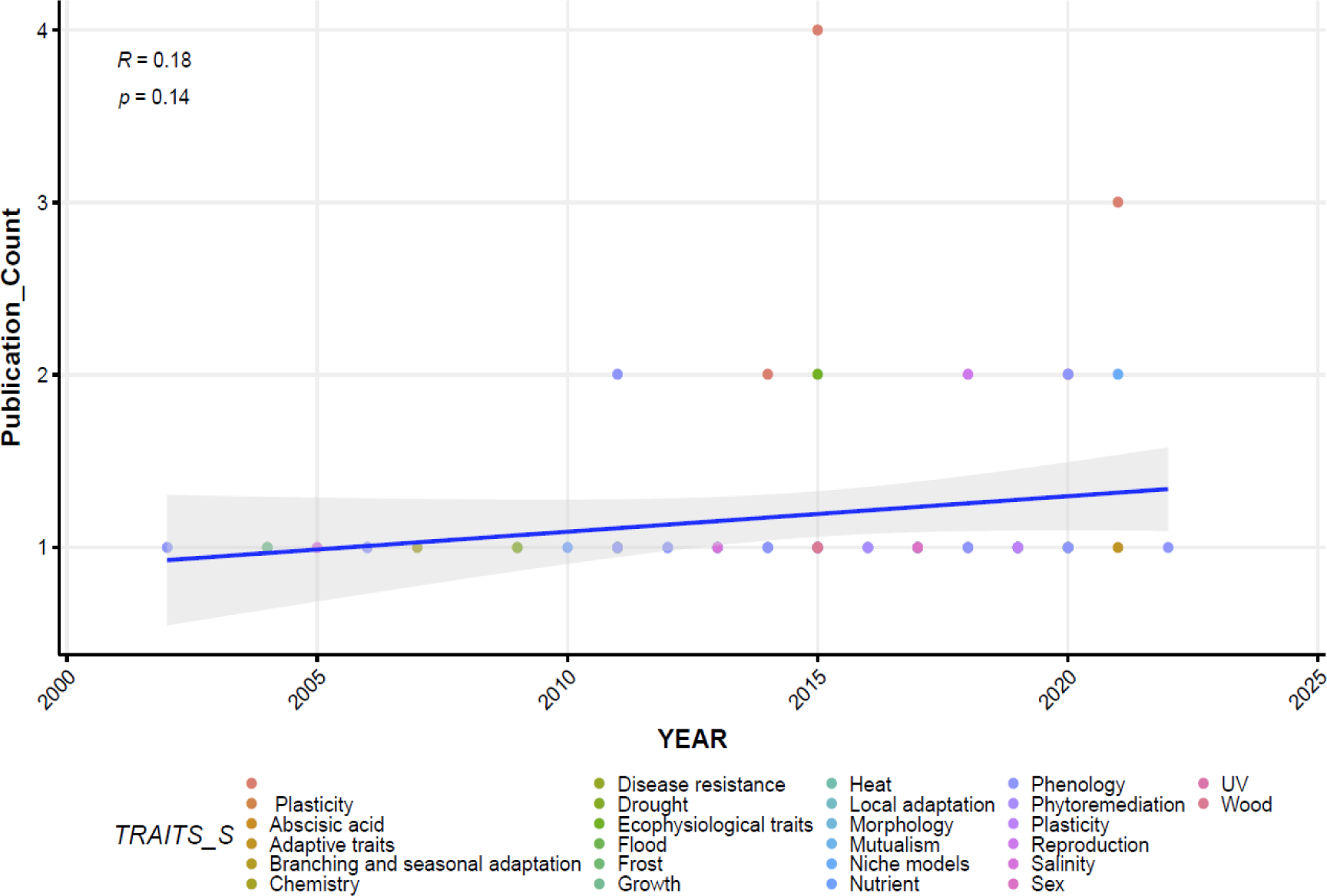
Temporal publications trend (incl. regression line) for broad spectra of adaptive traits in *Populus* spp. based on studies between yrs 2001 to 2021. Notes, the first dot (no name with color code) indicates not a specific trait study that can be categorized like the others.

**Table 1.** Literature sources of meta-analysis including the summary of species, main method, phenotypic traits of studies using *Populus* species.

| REFERENCES | REFERENCES | YEAR | TITLE | TRAITS_S | SPECIES |
| --- | --- | --- | --- | --- | --- |
| 1 | (Ding et al., 2020) | 2020 | Genetic parameters of growth and adaptive traits in aspen (*Populus tremuloides*): Implications for tree breeding in a warming world. | Phenology | *Populus tremuloides* |
| 2 | (Richards et al., 2020) | 2020 | Quantitative genetic architecture of adaptive phenology traits in the deciduous tree, *Populus trichocarpa* (Torr. and Gray). | Phenology | *Populus trichocarpa* |
| 3 | (Li et al., 2021a) | 2021 | Highly efficient C-to-T and A-to-G base editing in a *Populus* hybrid. |  | *Populus tremula* × *P. Alba* |
| 4 | (Welling et al., 2002) | 2002 | Independent activation of cold acclimation by low temperature and short photoperiod in hybrid aspen. | Phenology | Hybrid aspen |
| 5 | (Renaut et al., 2004) | 2004 | Responses of poplar to chilling temperatures: proteomic and physiological aspects. | Frost, Growth | Hybrid aspen |
| 6 | (Brosche et al., 2005) | 2005 | Gene expression and metabolite profiling of *Populus euphratica* growing in the Negev desert. | Salinity | *Populus euphratica* |
| 7 | (Bohlenius et al., 2006) | 2006 | CO/FT regulatory module controls timing of flowering and seasonal growth cessation in trees. | Phenology | *Populus tremula* |
| 8 | (Behnke et al., 2007) | 2007 | Transgenic, non-isoprene emitting poplars don't like it hot. | Chemistry | *Populus x canescens* |
| 9 | (Zhou et al., 2014) | 2014 | Exome resequencing reveals signatures of demographic and adaptive processes across the genome and range of black cottonwood (*Populus trichocarpa*). | - | *Populus trichocarpa* |
| 10 | (Yordanov et al., 2014) | 2014 | EARLY BUD-BREAK 1 (EBB1) is a regulator of release from seasonal dormancy in poplar trees. | - | *Populus tremula* × *P. Alba* |
| 11 | (Xiao et al., 2009) | 2009 | Physiological and proteomic responses of two contrasting *Populus cathayana* populations to drought stress. | Drought | *Populus cathayana* |
| 12 | (Wildhagen et al., 2010) | 2010 | Seasonal nitrogen cycling in the bark of field-grown Grey poplar is correlated with meteorological factors and gene expression of bark storage proteins. | Nutrient | *Populus tremula* × *P. Alba* |
| 13 | (Wildhagen et al., 2013) | 2012 | Low temperatures counteract short-day induced nitrogen storage, but not accumulation of bark storage protein transcripts in bark of grey poplar (*Populus x canescens*) trees. | Phenology, Nutrient | *Populus tremula* × *P. Alba* |
| 14 | (Rohde et al., 2011a) | 2011 | Temperature signals contribute to the timing of photoperiodic growth cessation and bud set in poplar. | Phenology | Hybrid poplar |
| 15 | (Qiu et al., 2011) | 2011 | Genome-scale transcriptome analysis of the desert poplar, *Populus euphratica*. | Drought | *Populus euphratica* |
| 16 | (Olson et al., 2013) | 2013 | The adaptive potential of *Populus balsamifera* L. to phenology requirements in a warmer global climate. | Phenology | *Populus balsamifera* |
| 17 | (Merino et al., 2014) | 2014 | Plantation forestry under global warming: hybrid poplars with improved thermotolerance provide new insights on the in vivo function of small heat shock protein chaperones. | Heat | *Populus tremula* × *P. Alba* |
| 18 | (Ma et al., 2013) | 2013 | Genomic insights into salt adaptation in a desert poplar. | Salinity | *Populus euphratica* |
| 19 | (Ko et al., 2011) | 2011 | Novel aspects of transcriptional regulation in the winter survival and maintenance mechanism of poplar. |  | *Populus trichocarpa* |
| 20 | (Keller et al., 2011) | 2011 | Climate-driven local adaptation of ecophysiology and phenology in balsam poplar, *Populus balsamifera* L. (*Salicaceae*) | Phenology, Ecophysiological traits | *Populus balsamifera* |
| 21 | (Hartikainen et al., 2009) | 2009 | "Emissions of volatile organic compounds and leaf structural characteristics of European aspen (*Populus tremula*) grown under elevated ozone and temperature. | Chemistry | *Populus tremula* |
| 22 | (Hall et al., 2011) | 2011 | Adaptive evolution of the *Populus tremula* photoperiod pathway. | Phenology | *Populus tremula* |
| 23 | (Ghelardini et al., 2014) | 2014 | Genetic architecture of spring and autumn phenology in *Salix*. | Phenology | *Populus trichocarpa* |
| 24 | (Fitzpatrick and Keller, 2015) | 2015 | Ecological genomics meets community-level modelling of biodiversity: mapping the genomic landscape of current and future environmental adaptation. | - | *Populus balsamifera* |
| 25 | (Fernandez-Martinez et al., 2013) | 2013 | Effect of environmental stress factors on ecophysiological traits and susceptibility to pathogens of five *Populus* clones throughout the growing season. | Ecophysiological traits | *Populus* |
| 26 | (Balint et al., 2013) | 2013 | Host genotype shapes the foliar fungal microbiome of balsam poplar (*Populus balsamifera*). | Mutualism | *Populus balsamifera* |
| 27 | (Azeez et al., 2014) | 2014 | A tree ortholog of APETALA1 mediates photoperiodic control of seasonal growth. | Phenology | Hybrid aspen |
| 28 | (Randriamanana et al., 2015) | 2015 | Interactive effects of supplemental UV-B and temperature in European aspen seedlings: Implications for growth, leaf traits, phenolic defense and associated organisms. | Ecophysiological traits, UV | *Populus tremula* |
| 29 | (Li et al., 2015) | 2015 | *Populus trichocarpa*. | - | *Populus trichocarpa* |
| 30 | (Zhang et al., 2020) | 2020 | Improved genome assembly provides new insights into genome evolution in a desert poplar (*Populus euphratica*). | - | *Populus euphratica* |
| 31 | (Zhang et al., 2019) | 2019 | Phenotypic and Genomic Local Adaptation across Latitude and Altitude in *Populus trichocarpa*. | Local adaptation | *Populus trichocarpa* |
| 32 | (Yu et al., 2019) | 2019 | Abscissic acid signalling mediates biomass trade-off and allocation in poplar. | Abscissic acid | Hybrid aspen |
| 33 | (Wulschleger et al., 2015) | 2015 | Genomics in a changing arctic: critical questions await the molecular ecologist. | - | *Populus* |
| 34 | (Wooley et al., 2020) | 2020 | Local adaptation and rapid evolution of aphids in response to genetic interactions with their cottonwood hosts. | Mutualism | *Populus angustifolia* |
| 35 | (Wojciechowska et al., 2020) | 2020 | Seasonal senescence of leaves and roots of *Populus trichocarpa*-is the scenario the same or different? | Phenology | *Populus trichocarpa* |
| 36 | (Wang et al., 2015) | 2015 | Evaluation of Appropriate Reference Genes for Reverse Transcription-Quantitative PCR Studies in Different Tissues of a Desert Poplar via Comparison of Different Algorithms. | - | *Populus euphratica* |
| 37 | (Vanden Broeck et al., 2018) | 2018 | Variability in DNA Methylation and Generational Plasticity in the Lombardy Poplar, a Single Genotype Worldwide Distributed Since the Eighteenth Century. | Plasticity | *Populus nigra* cv. *Italica Duroi* |
| 38 | (Samuilov et al., 2016) | 2016 | Lead uptake increases drought tolerance of wild type and transgenic poplar (*Populus tremula* x *P. alba*) overexpressing gsh 1. | Phytoremediation | *Populus tremula* × *P. Alba* |
| 39 | (Prunier et al., 2019) | 2019 | Gene copy number variations involved in balsam poplar (*Populus balsamifera* L.) adaptive variations. | Adaptive traits | *Populus balsamifera* |
| 40 | (Porth et al., 2015) | 2015 | Evolutionary Quantitative Genomics of *Populus trichocarpa*. | Growth, Phenology, Ecophysiological traits, Disease resistance, Wood | *Populus trichocarpa* |
| 41 | (Patil and Vijay, 2021) |  | Repetitive genomic regions and the inference of demographic history. | - | *Populus trichocarpa* |
| 42 | (Oubida et al., 2015) | 2015 | Partitioning of multivariate phenotypes using regression trees reveals complex patterns of adaptation to climate across the range of black cottonwood (*Populus trichocarpa*). | - | *Populus trichocarpa* |
| 43 | (Monson et al., 2020) | 2020 | High productivity in hybrid-poplar plantations without isoprene emission to the atmosphere. | Chemistry | Hybrid poplar |
| 44 | (Michelson et al., 2018) | 2018 | Autumn senescence in aspen is not triggered by day length. | Phenology | *Populus tremula* |
| 45 | (Miao et al., 2017) | 2017 | Sex-specific responses to winter flooding, spring waterlogging and post-flooding recovery in *Populus deltoides*. | Flood, Sex | *Populus deltoides* |
| 46 | (Menon et al., 2015) | 2015 | Population genetics of freeze tolerance among natural populations of *Populus balsamifera* across the growing season. | Phenology | *Populus balsamifera* |
| 47 | (Meirmans et al., 2017) | 2017 | History rather than hybridization determines population structure and adaptation in *Populus balsamifera*. | - | *Populus balsamifera* |
| 48 | (Maurya et al., 2020) | 2020 | A genetic framework for regulation and seasonal adaptation of shoot architecture in hybrid aspen. | Branching and seasonal adaptation | Hybrid aspen |
| 49 | (Liu and El-Kassaby, 2019) | 2019 | Phenotypic plasticity of natural *Populus trichocarpa* populations in response to temporally environmental change in a common garden. | Plasticity | *Populus trichocarpa* |
| 50 | (Li et al., 2021b) | 2021 | Genetic Architecture and Genome-Wide Adaptive Signatures Underlying Stem Lenticel Traits in *Populus tomentosa*. | - | *Populus tomentosa* |
| 51 | (Keller et al., 2017) | 2017 | Influence of Range Position on Locally Adaptive Gene-Environment Associations in *Populus* Flowering Time Genes. | Phenology | *Populus balsamifera* |
| 52 | (Holliday et al., 2016) | 2016 | Evidence for extensive parallelism but divergent genomic architecture of adaptation along altitudinal and latitudinal gradients in *Populus trichocarpa*. | Local adaptation | *Populus trichocarpa* |
| 53 | (Fitzpatrick et al., 2021) | 2021 | Experimental support for genomic prediction of climate maladaptation using the machine learning approach Gradient Forests. | - | *Populus balsamifera* |
| 54 | (Fahrenkrog et al., 2017) | 2017 | Population genomics of the eastern cottonwood (*Populus deltoides*). | Local adaptation | *Populus deltoides* |
| 55 | (Ding and Nilsson, 2016) | 2016 | Molecular regulation of phenology in trees — because the seasons they are a-changin | Phenology | *Populus tremula* |
| 56 | (DeWoody et al., 2015) | 2015 | Genetic and morphological differentiation in *Populus nigra* L.: isolation by colonization or isolation by adaptation? | Morphology | *Populus nigra* |
| 57 | (Dewan et al., 2018) | 2018 | Transgenerational effects in asexually reproduced offspring of *Populus*. | Reproduction | *Populus deltoides*<br>*Populus trichocarpa* |
| 58 | (Cooper et al., 2019) | 2019 | Genotypic variation in phenological plasticity: Reciprocal common gardens reveal adaptive responses to warmer springs but not to fall frost. | Frost, Phenology | *Populus fremontii* |
| 59 | (Chen et al., 2020) | 2020 | Survival in the Tropics despite isolation, inbreeding and asexual reproduction: insights from the genome of the world's southernmost poplar (*Populus ilicifolia*). | - | *Populus ilicifolia* |
| 60 | (McKown et al., 2018) | 2018 | Ecological genomics of variation in bud-break phenology and mechanisms of response to climate warming in *Populus trichocarpa* | Phenology | *Populus trichocarpa* |
| 62 | (Bothwell et al., 2021) | 2021 | Genetic data improves niche model discrimination and alters the direction and magnitude of climate change forecasts | Niche models | *Populus angustifolia*<br>*Populus fremontii* |
| 63 | (Apuli et al., 2021) | 2021 | The genetic basis of adaptation in phenology in an introduced population of Black Cottonwood<br>(*Populus trichocarpa*, Torr. &<br>Gray). | Adaptive traits | *Populus trichocarpa* |
| 64 | (Svystun et al., 2019) | 2019 | Modelling *Populus* autumn<br>phenology: The importance of<br>temperature and photoperiod | Phenology | *Populus tremula* |
| 65 | (Goessen et al., 2022) | 2022 | Coping with environmental<br>constraints: Geographically<br>divergent adaptive evolution and<br>germination plasticity in the<br>transcontinental *Populus*<br>*tremuloides* | Phenology | *Populus tremuloides* |

Future directions of genetic and genomic research application to improve adaptive traits in aspen are likely to include the following perspectives, 1) complex analytical approaches, more advanced and efficient sequencing and phenotyping methods as data volume and quality evolve exponentially; 2) regional and continental census and serial breeding, and testing of genetic and phenotypic variation among and within populations for tree improvement and conservation, e.g. assisted migration, tree breeding and selection, multivariate forestry, urban forestry applications; 3) genetic basis of phenotypic variation and environmental drivers of genetic adaptation, and clonal, familial, and population responses to stressors including climate change; 4) development of tools for GWAS and genomic selections for more accurate and rapid screening of clones and families for reforestation in terms of a multi-fold increase in genetic gains in commercial traits such as pulp yield and quality, fiber and wood quality, resistance against pests and pathogens; 5) growth and yield modeling considering genetic improvements in reforestation for commercial plantation; 6) development of pipelines for genetic, genomic and functional studies, sequencing tools, tree improvement that leverage research and development advances in both aspen and other non-model tree species; 7) latest genetic and biotechnology progress will facilitate the implementation of novel and advanced varieties of *Populus* trees, including *P. tremuloides*, for reforestation acceleration given the pipeline development from genetic engineering to seedling production and genetic improvement under local silvicultural settings. Finally, tree improvement networks, consortia and cooperations are promising regional and multi-regional approaches for research and sustainable operation to integrate the technological innovations and the extension efforts from the lab to endusers such as government and non-government agencies, landowners, and industrial stakeholders.

## Acknowledgements

The authors thank Dr Raju Soolanayakanahally (Agriculture and Agri-Food Canada), who continuously supported this study with valuable input.

## Notes

### Competing Interest Statement

The authors have declared no competing interest.

